# Dopaminergic neurons mediate male *Drosophila* courtship motivational state

**DOI:** 10.1101/733733

**Authors:** Guangxia Wang

## Abstract

Motivational states are important determinants of behavior. In *Drosophila melanogaster*, courtship behavior is robust and crucial for species continuation. However, the motivation of courtship behavior remains unexplored. We first find the phenomenon that courtship behavior is modulated by motivational state. A male fly courts another male fly when it first courts a decapitated female fly however, male– male courtship behavior rarely occurs under normal conditions. Male flies that have satisfied the need for sexual behavior show a decreased male–female sex drive. Therefore, in this phenomenon, the male fly’s courtship motivational state is induced by its exposure to female flies. Blocking dopaminergic neurons by expressing TNTe decreases motivational state-induced male–male courtship behavior without affecting male–female courtship behavior. Vision cues are another key component in sexually driven male–female courtship behavior. Here, we identify a base theory that the inner motivational state could eventually decide fly behavior.

## Introduction

Motivation provides behavior with purpose and intensity. A detailed neurobiological mechanism underlying state-dependent changes in behavior is lacking. To understand how the motivational state neural circuits are organized in the brain and how they impact neural circuits that direct behavior are major questions in neuroscience.

Studies on motivation in insects began with studying food-seeking behavior in the blowfly *Phormia regina* [1]. Although exposing gustatory receptor neurons on the proboscis to sugar always generated an electrophysiological response, the blowfly did not consistently respond by extending the proboscis. However, a food-deprived blowfly was more likely to respond with proboscis extension. Recently, a neural circuit that participates in the motivational control of appetitive memory was found. In fruit flies, appetitive memory expression is constrained by satiety and promoted by hunger. This group found that the stimulation of neurons that express neuropeptide F (dNPF) mimicked food deprivation and promoted memory performance in satiated flies. Robust appetitive memory performance requires the dNPF receptor expressed in dopaminergic neurons innervating a different region of the mushroom bodies. Blocking these dopaminergic neurons decreases memory performance in satiated flies, whereas stimulation suppresses memory performance in hungry flies [2].

Courtship behavior, as an innate behavior, has been widely studied since Seymour Benzer started the field of *Drosophila* neurogenetics at Caltech forty years ago. Each time, the male *Drosophila* performs the same ritual: orients towards a female, taps with its forelegs, sings a courtship song, licks the female’s genitalia and attempts copulation [3] after assessing the auditory, mechanosensory, visual and chemosensory signals from the target [4]. A female fly decides whether to accept a male fly based on the song he sings [5] and the pheromones he emits [6]. Neural circuits of courtship behavior are densely investigated based on the *fruitless* gene. Fruitless is a master regulator of sexuality in the fly [7] and was identified in 1963 by K.S. Gill and then cloned by Daisuke Yamamoto [8] and separately by the group of Hall, Bruce Bake and Barbara Taylor [9, 10]. Male courtship behavior can be induced in chromosomally female flies by expressing the male-specific isoform of *fru* in the female brain. This experiment demonstrated that this gene is key to male courtship behavior. Courtship circuitry based on the *fruitless* gene has been structurally mapped from sensory input to motor output [11], which provides the possibility of understanding how motivational state circuits direct motor output.

In this study, we build a paradigm to study the motivational state of male courtship. Male flies in this study first court decapitated female flies and are then exposed to male flies. Thus, we modulated the male fly’s sex motivation, increasing it to a high level. Sex-driven male–male courtship behavior was induced and compared with male flies without previous experience. No significant enhancement in male–female courtship was observed, which might be due to a ceiling effect. In contrast, sex-satisfied male flies show reduced courtship behavior to a new encounter with a female fly. Sex-driven male–male courtship behavior is primarily dependent on the vision cue input of the male fly. Blocking dopaminergic neuron transmission decreases sex-driven male–male courtship behavior. Whether increased dopamine levels could also rescue the low sex motivation level caused by satisfied courtship behavior needs to be explored.

## Results

### Male flies with high sexual motivation display state-dependent male–male courtship behavior, and sexually satisfied male flies show decreased sex drive to the next female fly

To investigate whether courtship behavior could be modulated by motivational state, we first changed the male fly’s sexual desire by exposing the male fly to a decapitated female fly. The 5-min period of time in which the male fly courted the decapitated female fly was called the exposure period (**FIG 1**), and the subsequent placing of the male fly with another male fly was called the test period (**FIG 1**), which was also 5 min. In the first exposure period, the male fly courted decapitated female flies with high intensity but was unable to copulate with the female flies, representing a high sexual drive state. In the control group, the male fly stayed alone without any sexual activity. In this situation, the male fly would court to the female fly vigorously without successful copulation because the decapitated female fly is not able to accept the male fly (**FIG 2 (A) (B) (C)**). Here, we use two parameters to describe the activity of courtship behavior. One is CI (courtship index), which means the percentage of total time the male fly spent on courtship during the observation period. The other is latency, which represents the time from the start of experiment to the time the male fly first displayed courtship behavior. Male–male courtship behavior is observed when male flies with a high sexual drive state and male flies without sexual behavior during the exposure period are immediately transferred to a different chamber and are exposed to other male flies (**FIG 2 (D) (E) (F)**). However, male flies rarely court male flies under normal conditions because this behavior is inhibited by the male pheromone 7-tricosone (7-T). Male flies display male–male courtship behavior mostly in the 1^st^ min and with short latency.

**FIG 1.**
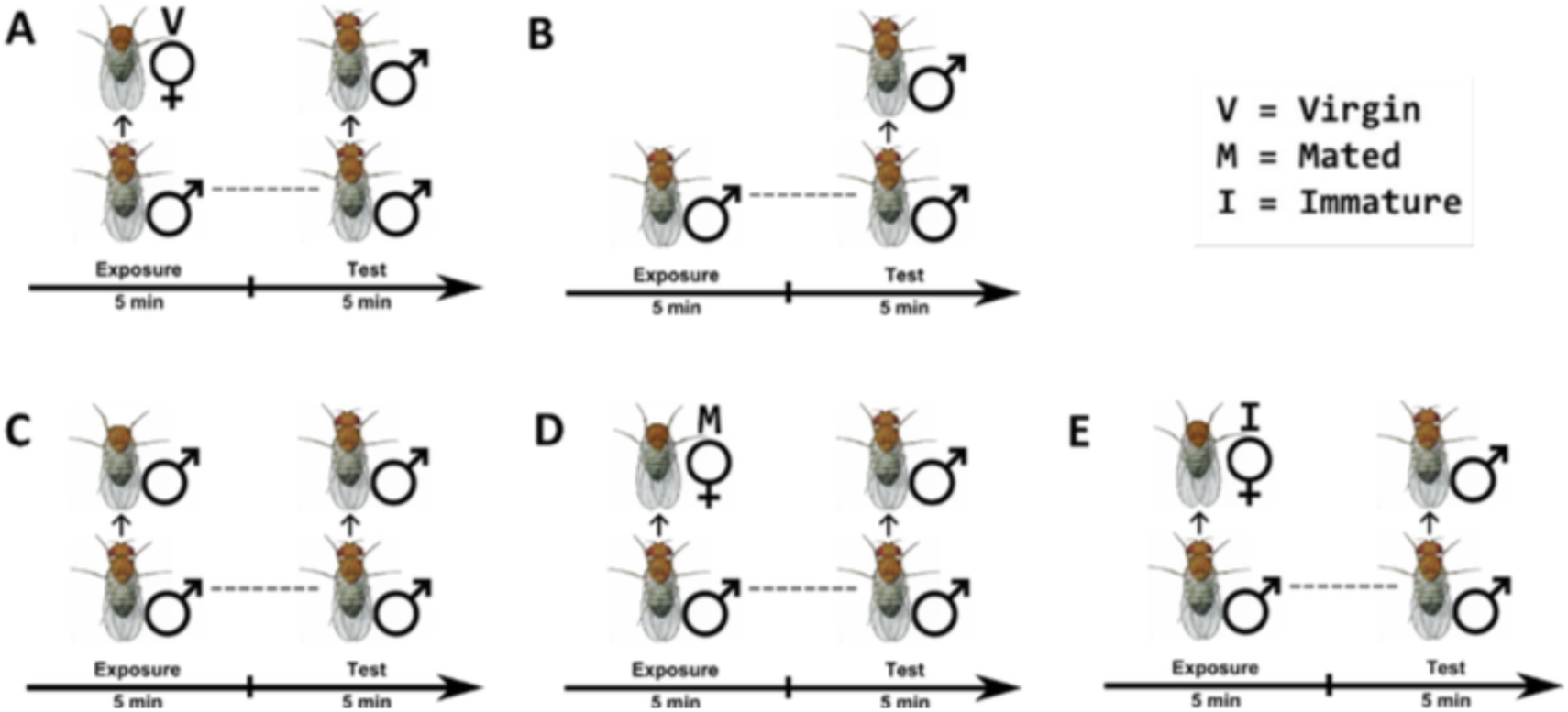
Paradigm for studying male *Drosophila* courtship motivation—two-round courtship behavior. During the exposure period, male flies are exposed to virgin female flies **(A)**, virgin male flies **(C)**, mated female flies **(D)**, and immature female flies **(E)** for 5 min each. Then, the male fly is transferred to another courtship chamber with a wing-cut male fly inside it for 5 min; this period is called the test period. **(B)**. Male flies stay alone during the exposure period for 5 min and then are transferred to another chamber with a wing-cut male fly inside it for 5 min. Paradigms (A) and (B) are mainly experimental and control groups, respectively.

**FIG 2.**
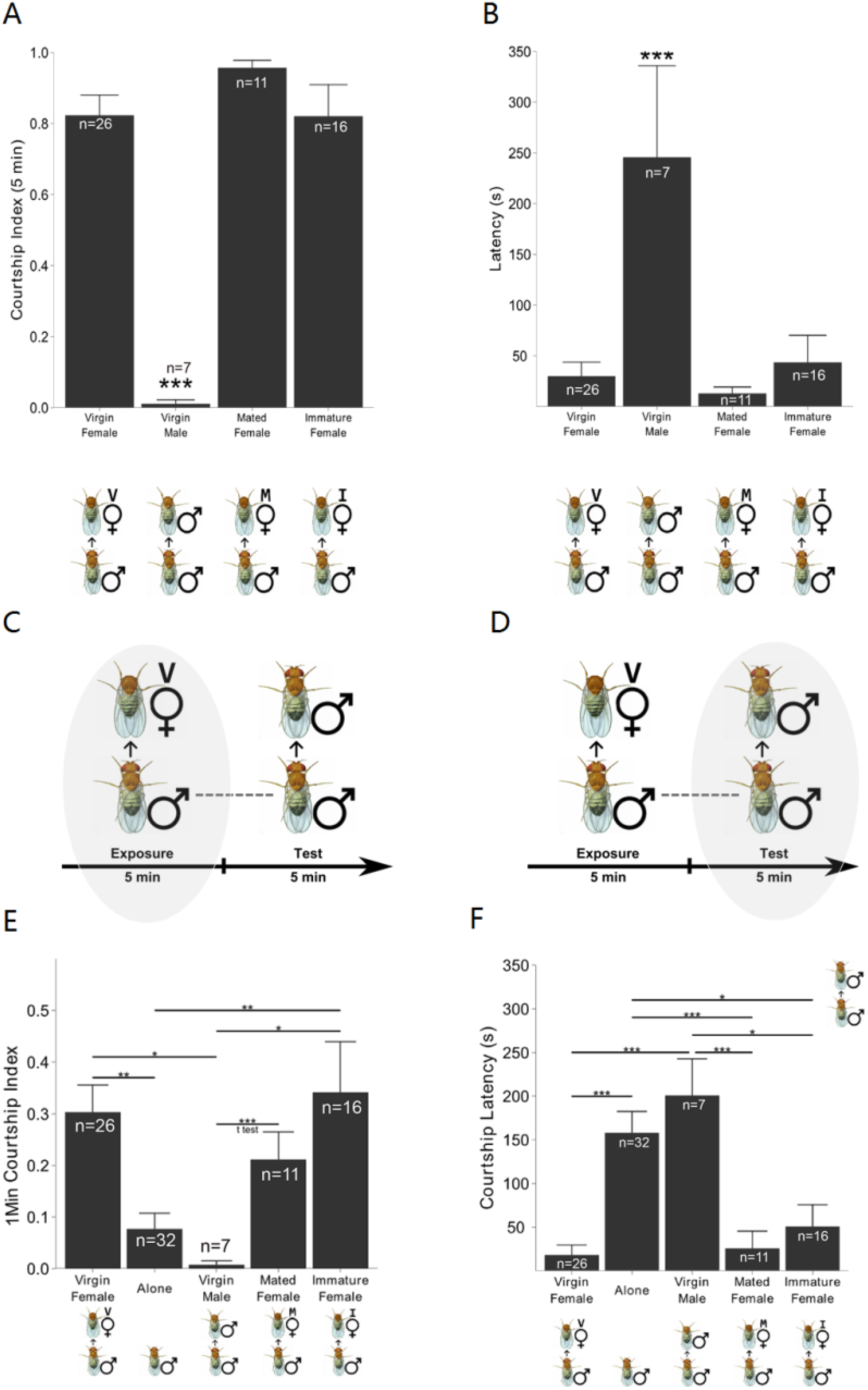
Male-male courtship behavior is induced after the male fly courts decapitated female flies intensely during the exposure period. **(A). (B)**. The wild-type male fly in the experimental group courts female flies intensely during the exposure period. **(A)**. Courtship index of male fly to female fly. **(B)**. Courtship latency of male fly to female fly. **(C)**. The experimental illustration indicates that the data for A and B come from the exposure period of the experimental group. **(D)**. The experimental illustration indicates that that E and F data display male–male courtship behavior in the test period. **(E)**. During the first minute of the test period, in the experimental group, the wild-type male fly displayed a significantly increased motivational male–male courtship index compared with that of the control group. **(F)**. During the first minute of the test period, in the experimental group, the wild-type male fly displayed a significantly decreased motivational courtship latency compared with that of the control group. n =7∼23. Mean ± s.e.m. The standard deviation is illustrated in the figure. ** indicates P<0.01, *** indicates P < 0.001.

We also tested whether sexually primed male flies display enhanced male–female courtship behavior (**FIG 3**). In contrast to the abovementioned experiment, this experiment paired male flies with intact virgin females instead of other male flies (**FIG 3 (A)**). We then observed male–female courtship behavior during both periods in different groups. Male flies did not show significantly enhanced male–female courtship behavior or significantly reduced latency. This result might be due to a ceiling effect of male–female courtship behavior during the exposure period.

**FIG 3.**
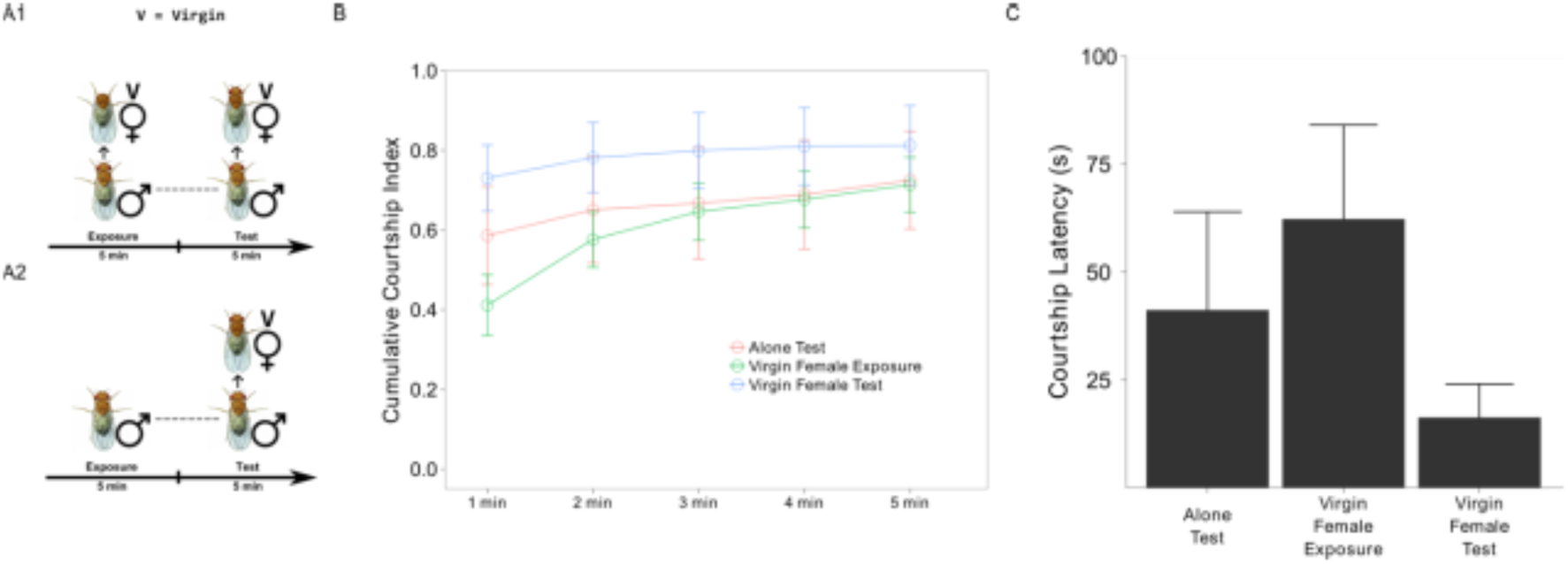
There is no increased male–female courtship behavior after the male fly intensely courts the first female fly during the test period. **(A1)**.**(A2)**. Behavior paradigm for the state-driven male–female courtship behavior. In this paradigm, the male fly is exposed to an intact female fly during the test period, which is the only difference from the previous paradigm. **(B)**. Male–female courtship index in the 1st min, 2nd min, 3rd min, 4th min, and 5th min of the exposure period and test period in both the experimental group and control group. There was no significant difference between these groups. **(C)**. Courtship latency in the exposure period and test period in both the experimental group and control group. There was no significant difference between these groups. n= 20. Mean ± s.e.m. The standard deviation is illustrated in the figure.

### Blocking TH neurons abolished motivational state-dependent male–male courtship behavior without decreasing male–female courtship behavior during the exposure period

Dopamine was shown to modulate male arousal and visual perception during heterosexual courtship [12, 13], locomotor activity [14], female sexual receptivity [15], male courtship conditioning [16], and ethanol-induced courtship inhibition [17]. Moreover, increased and decreased dopamine levels induce homosexuality in *Drosophila* [18, 19]. Therefore, we tested the function of dopamine in this experiment. Using TH-gal4 driving TNT expression, we blocked TH neurotransmission. We found that male flies did not display enhanced male–male courtship behavior compared with control male flies (**FIG 4 (A) (B) (C)**), while male–female courtship behavior remained unaffected during the exposure period (**FIG 4 (D) (E) (F)**). Experiments with Gal80ts expression in the fly could exclude developmental effects. This motivational state-dependent behavior could be transient; thus, we hypothesize that the developmental effect of TNTe might be slight. As a control group, we blocked the activity of octopaminergic neurons during the test period, and this manipulation did not influence the courtship motivational state during the test period (**FIG 5**).

**FIG 4.**
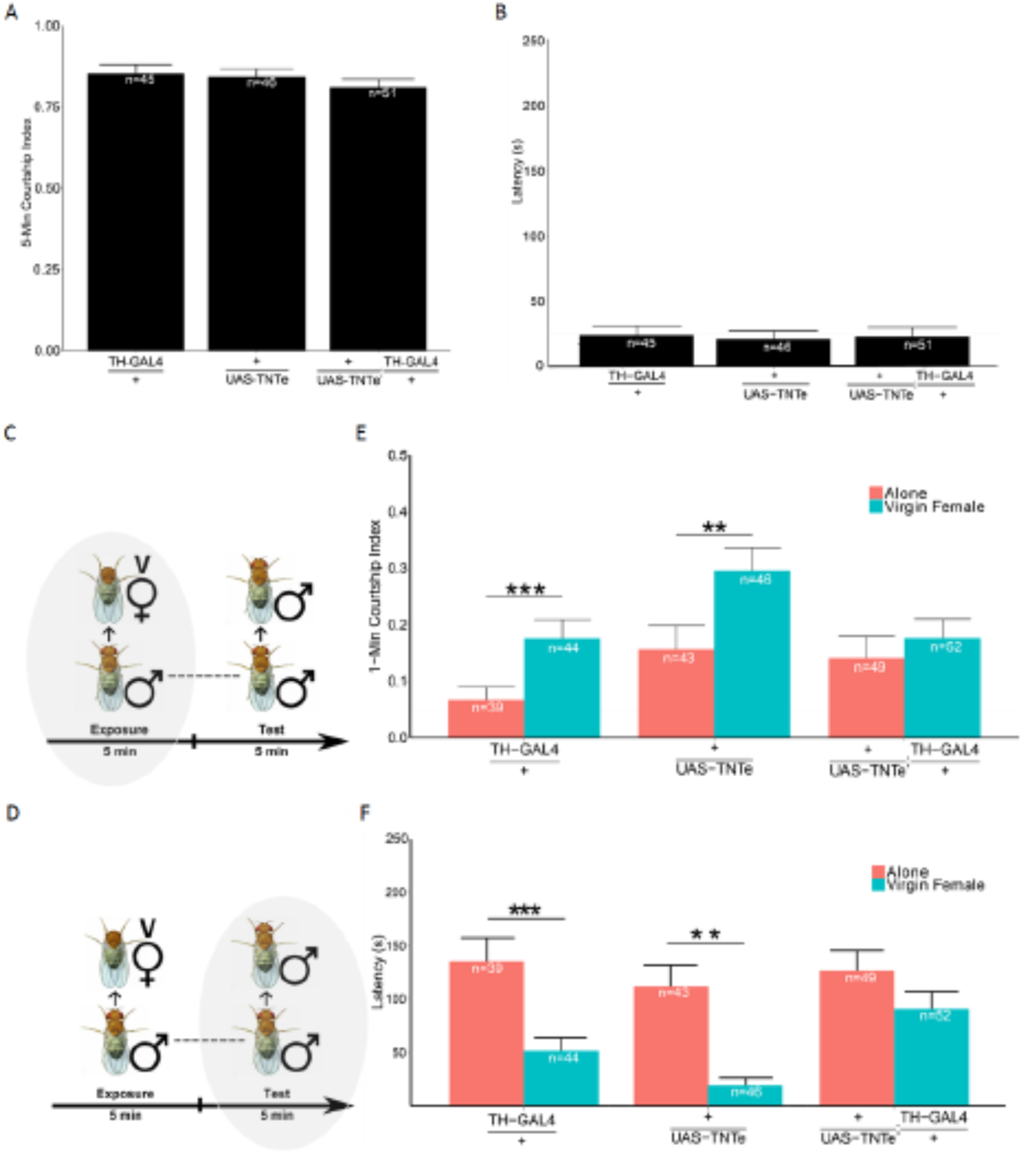
Blocking dopaminergic neurotransmission destroys sex-driven male-male courtship behavior, but it does not influence male–female courtship behavior during the exposure period. **(A), (B)**. During the exposure period, male flies that block dopaminergic transmission (UAS-TNTe/+; TH-GAL4/+) display the same intense male– female courtship behavior as the control fly, TH-Gal4/+ and UAS-TNTe/+. **(A)**. Courtship index for male–female courtship behavior. **(B)**. Courtship latency for male–female courtship behavior. **(C)**. Data for A and B come from male–female courtship behavior during the exposure period, n=45∼51. **(D)**. Data for A and B come from male–male courtship behavior during the test period. **(E), (F)**. During the test period, male flies (UAS-TNTe/+; TH-GAL4/+) that block dopaminergic transmission display no increased male– male courtship behavior compared with the control group. Control fly TH-Gal4/+ and UAS-TNTe/+ display increased sex-driven male–male courtship behavior. **(E)**. Courtship index of male–male courtship behavior. For the UAS-TNTe/+genotype, p = 0.0003148 for the TH-GAL4/+ genotype, p = 0.00342, **(F)**. Courtship latency of male–male courtship behavior. For the UAS-TNTe/+ genotype, p = 3.637e-05, for the TH-GAL4/+ genotype, p = 0.007924. n =39∼52. The standard deviation is illustrated in the figure. Mean ± s.e.m. * indicates P<0.05, ** indicates that P < 0.01.

**FIG 5.**
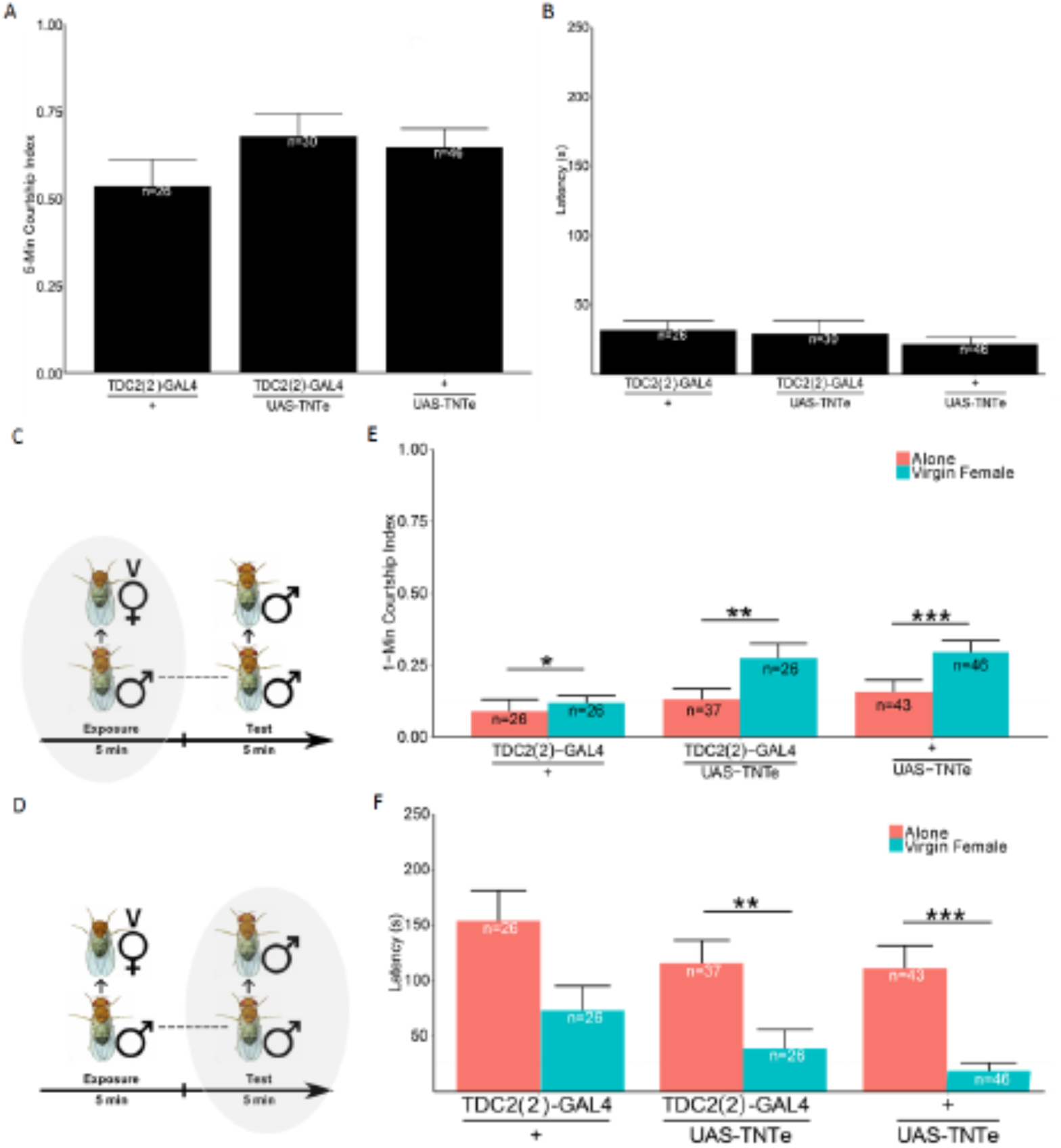
Blocking octopaminergic neurotransmission has no influence on sex-driven male–male courtship behavior, and it does not influence male–female courtship behavior during the exposure period. **(A), (B)**. During the exposure period, male flies that block dopaminergic transmission (UAS-TNTe/+; TDC2(2)-GAL4/+) display the same intense male–female courtship behavior as CS female flies with control flies, TDC2(2)-Gal4/+ and UAS-TNTe/+. **(A)**. Courtship index for male–female courtship behavior. **(B)**. Courtship latency for male–female courtship behavior. **(C)**. Data for A and B come from male–female courtship behavior during the exposure period, n = 26∼46. **(D)**. Data for A and B come from male–male courtship behavior during the test period. **(E), (F)**. During the test period, male flies (UAS-TNTe/+; TDC2(2)-GAL4/+) that block dopaminergic transmission display increased male–male courtship behavior compared with the control group. Additionally, control fly TDC2(2)-Gal4/+ and UAS-TNTe/+ display increased sex-driven male–male courtship behavior. **(E)**. Courtship index of male–male courtship behavior. **(F)**. Courtship latency of male–male courtship behavior, n = 26∼46. The standard deviation is illustrated in the figure. Mean ± s.e.m. * indicates P<0.05, ** indicates that P < 0.01.

### Sex-driven male–male courtship behavior depends mostly on vision perception

In addition to the inner state, sensory stimuli could also induce courtship behavior. Therefore, we want to find the main sensory modality that could play a role in sex-driven male–male courtship behavior. This male–male courtship behavior could start even with a long distance existing between the two flies. We focus on olfaction and vision first. Male flies with defective olfaction and those with white eyes (*W*^*1118*^ and WCS; **FIG 6 (C) (D)** and **FIG 6 (E) (F)**, respectively) were separately tested. We found that male flies with vision defects could not produce this kind of behavior (**FIG 6 (A) (B)**), while flies with olfaction defects showed no difference. Immobile target flies without any pheromones were used in the test period (**FIG 6**).

**FIG 6.**
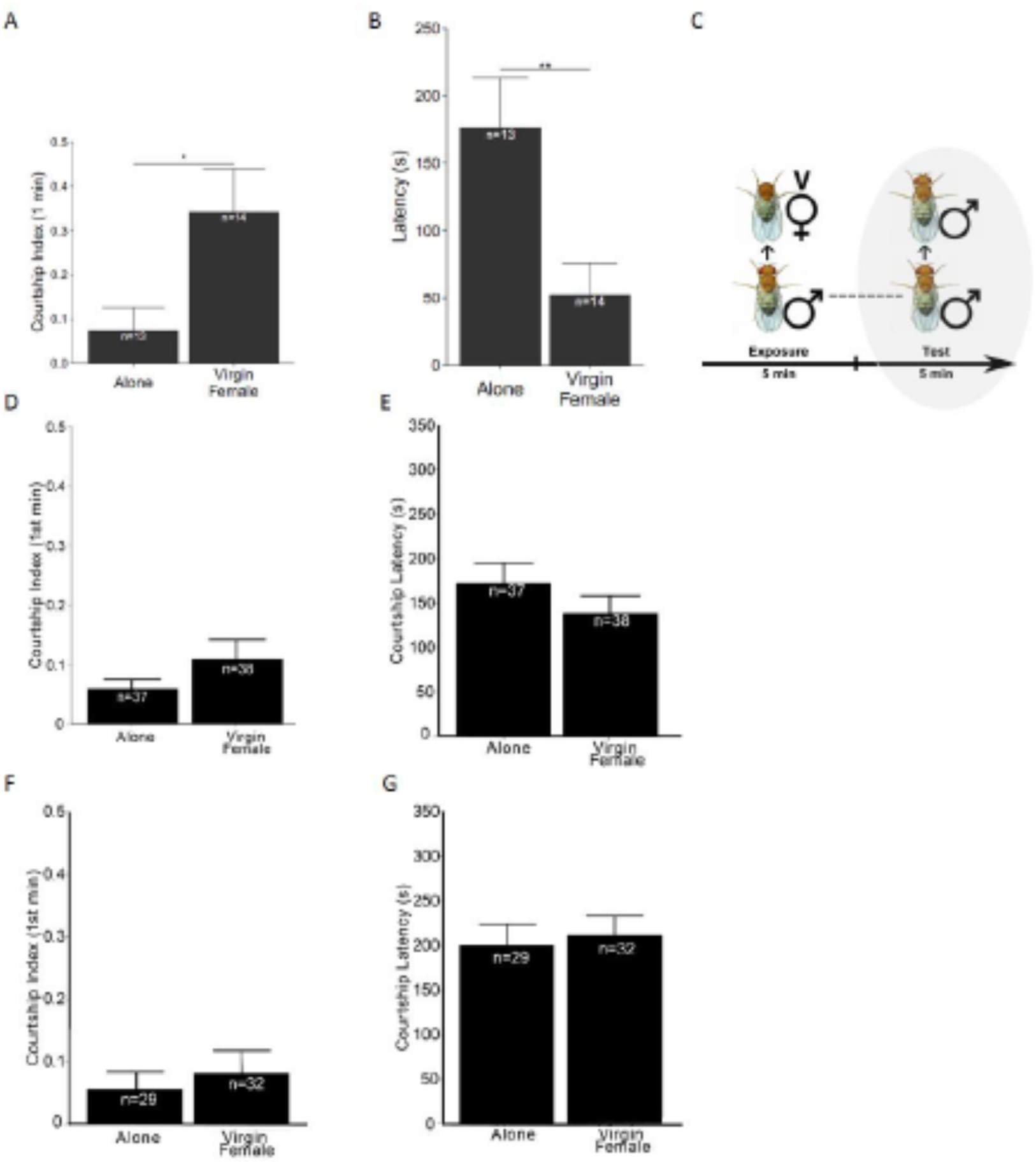
Vision deprivation but not olfaction deprivation decreases sex-driven male– male courtship behavior. **(A). (B)**. After olfaction deprivation, the wild-type male fly could still display sex-driven male–male courtship behavior. **(A)**. Male–male courtship index (P=0.0009536, Wilcoxon rank sum test), **(B)**. Male–male courtship latency. (P=0.008138, Wilcoxon rank sum test), n =13∼14. **(C)**. The data of (A), (B), (D), (E), (F), (G) come from male–male courtship behavior during the exposure period in the experimental group, n=45∼51. *W*^*1118*^ and WCS are used as the test flies with vision deprivation in (D), (E) and (F), (G). They do not display sex-driven male–male courtship behavior. **(D)** and **(F)**, Courtship index. **(E)** and **(G)**, Courtship latency. n=34∼37. Mean ± s.e.m. The standard deviation is illustrated in the figure. * indicates P < 0.05, **indicates that P < 0.01.

## Discussion

Sensory stimuli play important roles in animal output behavior, but the inner state of the animal decides the output behavior. The inner state enables animals to behave flexibly and will help the animal satisfy its needs to the maximum extent. Courtship behavior is important for species continuation, and the inner state that underlies this behavior is crucial for animal reproduction. The fixed style of courtship behavior and relatively clear neural circuit based on the courtship genes *doublesex* and *fru* have attracted the attention of many researchers [11, 20-24]. However, the neural circuit that underlies the inner state of courtship behavior remains unclear. Our research has filled in the blanks.

The question of whether internal state influences courtship behavior can be broken down into three questions: 1. whether the inner state of *Drosophila* influences its output behavior; 2. which neural circuit underlies the inner state and how it codes and influences courtship behavior; and, 3. how the neural mechanisms interact with the decision center, motor center and sensory inputs to decide the influence of the inner state on output behavior. To date, we have answered the first two questions, and the third question has not been solved in our research.

For the first question, we ask whether the inner state of *Drosophila* will influence the output of *Drosophila* courtship behavior. First, we needed to change the inner state of *Drosophila*. Through a two-round courtship behavior paradigm, we successfully raised the inner state in *Drosophila* courtship behavior. The difficulty and novelty of this part of the work was to display this raised courtship inner state. Here, male–male courtship behavior provided a good standard by which to display the raised inner courtship state. Under normal conditions, male–male courtship behavior rarely occurs. In addition, male–male courtship behavior will happen when *Drosophila* have an increased internal courtship state and can rescue the ceiling effect of male–female courtship behavior.

In contrast, a decreased inner courtship state is displayed during male–female courtship behavior after successful copulation with a previous female fly. This idea is similar to that of a study in which the male fly’s courtship state decreased after multiple copulations with a female fly [25]. These two methods can successfully adjust the inner state of *Drosophila*. At the same time, *Drosophila*’s courtship behavior changed as its inner state changed.

For the second question, we ask which neural mechanisms underlie the *Drosophila* inner state. In mammals, including humans, several studies have proven that the dopamine system is important in adjusting the state of behavior. For example, patients who take L-DOPA have increased sexual activity. In *Drosophila*, some studies have proven that the dopamine system takes part in modulating sexual behavior. Administering dopamine receptor agonists to *Drosophila* increases male–female courtship behavior [26]. These studies have shown the role of dopamine in modulating courtship behavior. However, this pharmacology study shows that these agonists have an unknown effect on behavior. Therefore, we built a paradigm to study this result in-depth via behavior and genetic methods.

In our research, we found that the activity of dopamine neurons will display the inner state of courtship behavior. Blocking dopaminergic neuron transmission via expressing UAS-TNTe driven by TH-Gal_4_ will decrease the raised inner state-induced male–male courtship behavior. Therefore, we ask whether increased male–male courtship behavior will increase TH release. To answer these questions, specific subtypes of TH neurons needed to be identified first, which depends on a genetic method.

In *Drosophila melanogaster*, TH neurons overlap with cells in *fru* circuits, including aSP4 and aSP13. In addition, aSP4 neurons’ presynaptic area projects to the postsynaptic area of P1, which is the commander of courtship circuit neurons, SMPa [11]. aSP4 and PPL2ab dopaminergic neurons are important candidates in the inner state of courtship behavior. aSP4 dopaminergic neurons belong to the PAL subtype. In male flies, courtship behavior will be abolished if these neurons are feminized [27]. The overexpression of tyrosine hydroxylase could increase courtship intensity in old male flies [28].

In addition, one strategy is labeling the neurons activated by the increased inner courtship state of *Drosophila*. Here, we tried two methods. One method uses the DopR-Tango system, which would express the LexA promoter when DopR1 is activated. Then, we produced a cross with LexAop-reporter protein to ensure the expression of dopamine-dependent fluorescence [29].

However, because the mutant induced by gene insertion caused incomplete wings, the system did not suit our courtship behavior. The dopamine receptor only includes DopR, and not studying all receptors is a limitation of this research. If we can answer these two problems, it will provide a good system for the research.

With another method, we identified the neurons activated by increased inner courtship state via the CaMPARI system [30], which is a calcium binding protein that when irreversibly combined with calcium, changes its color from green to red under ultraviolet light. We have found some activity-labeled neurons via this method. This result needs further study. In addition, red light could replace ultraviolet rays, which would be more suitable for flies because they cannot sense red light.

For the third question, we asked how dopamine neurons changed the inner state. In *Drosophila* courtship behavior, some *fru*-positive neurons project to *Drosophila* sensory integration and behavior output centers. P1 interneurons are important for triggering courtship behavior. They receive sexual sensory information from female [31, 32] flies and project to motor neurons [32, 33]. The activation of P1 neurons alone will induce male courtship behavior toward a fake female such as one made of rubber [34]. It remained unknown whether dopamine neurons could directly influence motor output neurons to induce male–male courtship behavior and via which dopamine receptor.

Our research found that after the male fly raised the inner courtship state, the vision cue played an important role in male–male courtship behavior. Whether dopamine neurons sensitize visual inputs or whether they utilize the visual cue as a dependent cue remains to be explored. This answer depends on the detailed relationship between vision input circuits and dopamine neurons.

Sensory neurons help male flies identify the target, while the center brain’s state modulates *Drosophila* output behavior. The relationship between dopamine neurons and vision sensory circuits remains unknown. Dopamine neurons might play a dependent role in determining courtship motivational state behavior, or the inner state might sensitize the vision input circuit. The specific TH subtype that modulates inner state-induced male–male courtship behavior remains unknown. Whether increased dopamine levels will rescue vision deprivation-induced motivational state-induced male–male courtship deficiency remains unknown. These questions require further study.

## Author Contributions

Guangxia Wang designed, performed and analyzed all the behavior experiments and wrote the manuscript. Aike Guo helped discuss the experimental design. There is no conflict of interest in this paper.

## Acknowledgements

Bangyu Zhou wrote the Video imprinter software for behavior analysis, which works well, especially on Windows systems. Tong Liu assisted in discussing the work. Ying Wang assisted with the fly preparation. This work was supported by the 973 Program Grant 2011CBA00400 (to A.G.), Natural Science Foundation of China Grants 30921064, 90820008, and 31130027 (to A.G.), and the Strategic Priority Research Program of Chinese Academy of Science, Grant No.XDB32010100.

## Methods

### Fly strains

The fly strains used in these experiments contain CS, *W*^*1118*^ and WCS, UAS-TNTe, and TH-Gal4 [13]. All flies are raised on Bloomington standard food. Then, they are placed in 23∼25°C, 50∼70% humidity, 12 h light/12 h dark rooms and incubators. All female flies and male flies in the following experiments are virgin, and they are picked up in the condition of CO_2_ anaesthetization. After that, female virgin flies are raised as a group of 10, and virgin male flies are raised singly in small tubes; potassium sorbate replaces propionic acid as a food preservative. Male flies are 4∼8 days old, and female flies are 4∼10 days old in this study. All target flies are wild CS. They are both first generation crosses, unless otherwise stated.

### Background washing

Virgin females flies in the experiments (TH-GaL4, TDC2(2)-Gal4, UAS-TNTe [14]) are crossed with male flies (*W*^*1118*^), then the next generation of virgin female flies is picked and crossed with male flies (*W*^*1118*^) for six generations, after which we crossed the flies with a double balancer to obtain *Drosophila melanogaster* with pure, identical genetic backgrounds.

All flies are virgin unless otherwise stated. Male flies were raised as single flies, and female flies were raised in groups of 10 flies. All the target flies are CS.

### Behavior experiment

All experiments were performed in a 4-layer courtship chamber and in a custom transparent cube (approximately 100 cm x 50 cm x 80 cm) to ensure constant temperature and humidity. During the experiment, the temperature was maintained at 23°C and 60% humidity. Inside the chamber, wide-angle earthquake-proof DVs (SONY) were fixed to record the entire experiment, and after that, we analyzed the video data with a self-coding video imprinter. During the experiment, we designed a double-blind method to exclude subjective deviations in the results. The procedures of the experiments are as follows: The target male fly and female fly were placed under frozen anaesthetization and then transferred to a temperature-adjustable 4°C copper plate. The female fly is decapitated, as the target fly during the exposure period, to ensure the induction of sexual desire, and the male is wing-cut via fine scissors, as is the target fly, during the test period keep this fly and the male test fly separated. The flies are placed in adjacent, but different, layers of the courtship chamber. The active test male fly freely moved into one layer of the chamber, next to the female fly.

During the exposure period, after the female fly recovered, it was transferred into the female fly’s chamber for 5 min, and then, quickly and softly, the male fly was moved into another chamber with a wing-cut male fly inside it for 5 min, which was the test period. In the control group, during the first period, the male fly is transferred into an empty chamber for 5 min. In the negative control experiment, during the exposure period, the male fly is exposed to a decapitated male fly. During the test period, they were both exposed to normal wing-cut male flies.

In the experiment for desire satisfaction, we placed a male fly and two normal virgin females in 3 different layers of the courtship chamber. In addition, we placed the male with the first female, and after successful copulation with this first female fly, the male fly was immediately transferred to a chamber with a second virgin female fly for the second courtship behavior. Then, the courtship index and latency for the first 5 min of the two courtship behaviors were separately calculated to indicate the strength of courtship desire.

### Olfaction deprivation experiment

Flies were placed on a 4°C copper plate after frozen anaesthetization, and then, the antenna and maxillary palp were removed with tweezers. After that, the male fly was transferred back into its tube and was allowed to recover for 2 days before we performed the experiment.

### Data analysis

Videos recorded were analyzed by self-coding video imprinter software running specifically under a Windows system. We tracked each step of the courtship behavior, including orientation, following, singing courtship song, licking female genitalia, attempting copulation, and copulation. Then, we analyzed the script of tracking data and drew pictures via R code. The Courtship Index is defined as the percentage of the occurrence time of courtship behavior to the total observation time. Courtship latency is defined as the time from the beginning of the experiment to the initiation of courtship behavior. Because the data are not normal distribution, the Wilcoxon rank test was used for significance difference detection. Adobe Illustrator was used as the phototypesetting software.

